# Dynamic nuclear polarization illuminates key protein-lipid interactions in the native bacterial cell envelope

**DOI:** 10.1101/2023.05.18.541325

**Authors:** James E. Kent, Bryce E. Ackermann, Galia T. Debelouchina, Francesca M. Marassi

## Abstract

Elucidating the structure and interactions of proteins in native environments has become a fundamental goal of structural biology. Nuclear magnetic resonance (NMR) spectroscopy is well suited for this task but often suffers from low sensitivity, especially in complex biological settings. Here, we use a sensitivity-enhancement technique called dynamic nuclear polarization (DNP) to overcome this challenge. We apply DNP to capture the membrane interactions of the outer membrane protein Ail, a key component of the host invasion pathway of *Yersinia pestis*. We show that the DNP-enhanced NMR spectra of Ail in native bacterial cell envelopes are well resolved and enriched in correlations that are invisible in conventional solid-state NMR experiments. Furthermore, we demonstrate the ability of DNP to capture elusive interactions between the protein and the surrounding lipopolysaccharide layer. Our results support a model where the extracellular loop arginine residues remodel the membrane environment, a process that is crucial for host invasion and pathogenesis.

---

Understanding how proteins interact with their environments at the atomic level is critical for gaining mechanistic biomedical insights. Nuclear magnetic resonance (NMR) is exceptionally well suited for this purpose as NMR signals are highly susceptible to the local environment of their corresponding sites, and therefore, capable of reporting on even very weak intermolecular interactions. Taking advantage of these capabilities, the growing fields of *in situ* and *in cell* NMR *(1, 2)* present new opportunities for examining protein structure and interactions in native biological contexts, including the investigation of protein aggregation in cellular lysates *(3)*, the characterization of protein function in native membranes *(4-6)*, and the description of protein-drug interactions in cells *(7)*. Despite these advances, however, a known limitation of NMR is its inherently low signal intensity. The extensive signal averaging times required to accumulate sufficient signal over noise can be impractical for complex samples such as native membranes and cells, and can compromise sample stability. Moreover, signals from sites undergoing chemical or dynamic exchange on the millisecond time scale are difficult to detect. These problems can be particularly limiting for solid-state NMR spectroscopy, which still predominantly relies on the observation of low sensitivity nuclei such as ^13^C and ^15^N, and can be sensitive to molecular motions that interfere with the recoupling and decoupling schemes employed in the experiments *(8, 9)*.

The development of high field Dynamic Nuclear Polarization (DNP) for solid-state NMR spectroscopy *(10)*, reviewed in ref. *(11-13)*, provides a key avenue for overcoming these challenges. In a DNP experiment, polarization is transferred from unpaired electron spins to nuclei, typically resulting in enhancements of 10–150 for biological samples. To this end, samples are doped with stable biradicals, and experiments are performed at 90–100 K under magic angle spinning (MAS) conditions. DNP has been used to investigate a wide range of systems, including amyloid fibrils *(3, 14)*, nucleic acids and chromatin polymers *(15-17)*, membrane-embedded proteins *(4, 18, 19)*, and intact cells *(20-22)*. Here, we use DNP-enhanced solid-state NMR spectroscopy to investigate the lipid interactions of the protein Ail, a virulence factor of the plague pathogen *Yersinia pestis*, in intact bacterial cell envelopes.

Previously, we showed that the *Yersinia pestis* surface protein Ail can be expressed in the outer membrane of *E. coli* cells for *in situ* solid-state NMR structural studies at atomic-resolution *(5)*. The solid-state NMR spectra from isolated bacterial cell envelopes (including both inner and outer membranes, and interleaving peptidoglycan layer) reflect the eight-stranded β-barrel structure of Ail, and reveal sites that interact with bacterial outer membrane components and human serum factors. Moreover, since Ail expression confers some of its key virulence phenotypes to *E. coli*, the structural data from these samples can be correlated with activity. Our previous NMR studies of Ail with bacterial cell envelopes, lipid nanodiscs, and liposomes, together with molecular dynamics (MD) simulations and microbiology assays, all point to specific interactions between key basic sidechains of Ail with the lipopolysaccharide (LPS) lipids of the bacterial outer membrane *(5, 23)*. These interactions are important for virulence as in *Y. pestis* bacteria, Ail and LPS have co-evolved to confer resistance to human innate immunity, and act cooperatively to enhance pathogen survival in serum, antibiotic resistance, and cell envelope integrity *(24-27)*. Nevertheless, the NMR signals from the LPS binding sites of Ail have been challenging to detect and this limits our understanding of the precise molecular mechanism for this interaction. In this study, we show that the DNP-enhanced solid-state NMR spectra of Ail in bacterial membranes provide direct evidence of contacts between key Arg sidechains of Ail and the outer membrane LPS.

To examine the potential of enhancing the NMR signals from Ail, *in situ*, using DNP, we used a ^15^N/^13^C labeled Ail sample in intact *E. coli* cell envelopes. The sample was prepared, as previously described *(5)*, by first growing cells in unlabeled media and then transferring them to isotopically labeled media supplemented with both rifampicin and IPTG. Rifampicin halts transcription of the *E. coli* genome and suppresses endogenous protein production, while IPTG induces Ail expression *(28)*. This protocol of targeted isotopic labeling is critical for suppressing background NMR signals from other cellular components and for yielding high-resolution solid-state NMR spectra of Ail *in situ (5)*. We then doped the sample with 10 mM of the nitroxide-based biradical AMUPol, and cryoprotected the sample in a mixture of 60% glycerol-d_8_, 35% D_2_O, and 5% H_2_O. DNP experiments were performed at 600 MHz and 100 K, with 12 kHz MAS frequency.

The ^15^N-filtered two-dimensional ^15^N/^13^C correlation NCA spectrum acquired with DNP at 100K also exhibits significant signal enhancement. It compares very favorably with its counterpart acquired at 278K, notwithstanding the line broadening, with numerous identifiable signals that could be assigned based on their 278K assignments (Fig. 2). Twenty Gly peaks are expected based on the amino acid sequence of Ail and additional signals are observed in the ^15^N/^13^CA spectral region associated with Gly (^13^C: 42-48 ppm; ^15^N: 101-121 ppm), with one signal (^13^C: 44.5 ppm; ^15^N: 121.0 ppm) tentatively assignable to Gly19 in the first extracellular loop of Ail, based on comparison with resonance assignments made in decylphosphocholine micelles *(29-31)*. These results illustrate the greater amount of information that is available from NMR spectra obtained at low temperature. Moreover, the observation of discrete, resolvable peaks reflects relative order, within narrow conformational ensembles, for their corresponding protein sites rather than high conformational disorder.

**Figure 1.**
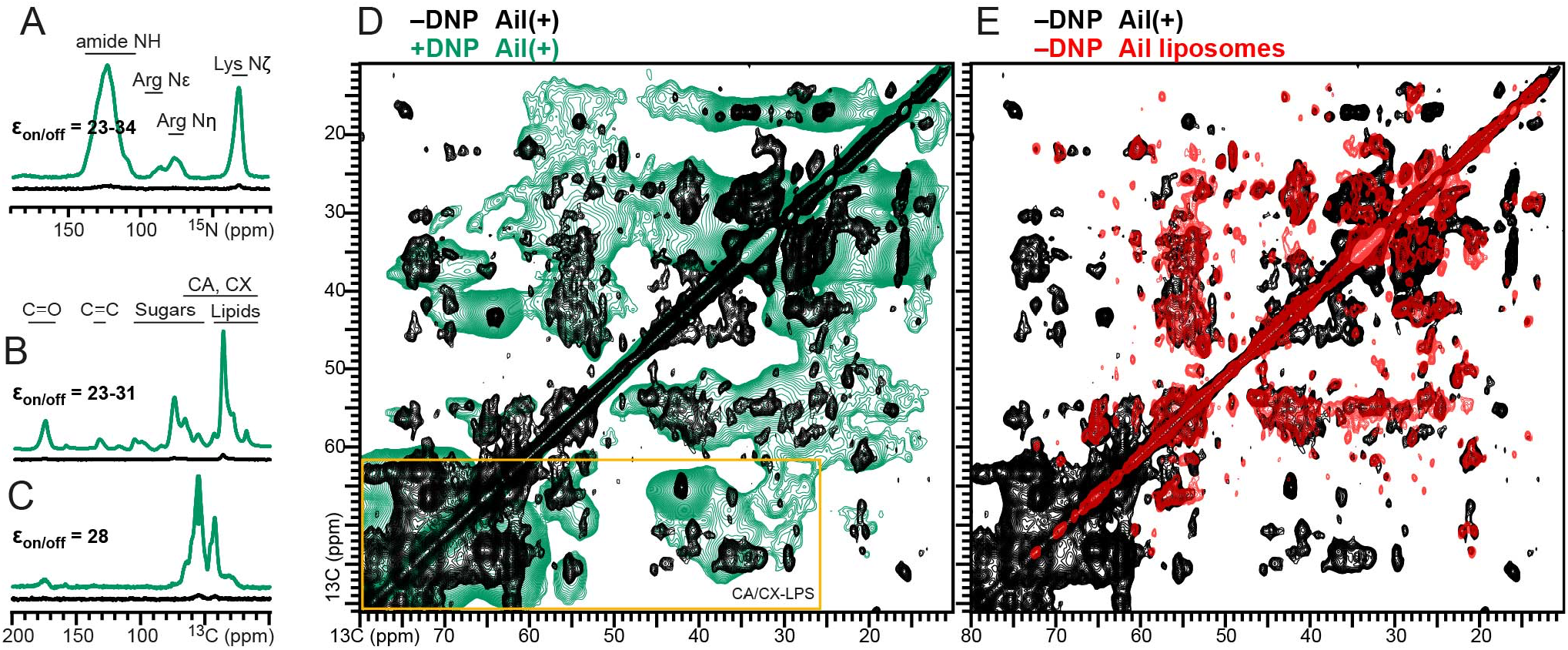
DNP signal enhancement of the ^15^N and ^13^C MAS solid-state NMR spectra of Ail in *E. coli* cell envelopes. **(A-C)** One dimensional spectra acquired with (A) ^1^H-^15^N CP, (B) ^1^H-^13^C CP, or (C) ^1^H-^15^N-^13^C double CP, and with (green) or without (black) microwave irradiation. Signal enhancement factors (ε_on/off_) were measured as the ratio of signal intensity observed with and without microwave irradiation. **(D, E)** Two dimensional ^13^C/^13^C correlation spectrum of Ail in *E. coli* cell envelopes acquired with DNP (green). The spectra from Ail in *E. coli* cell envelopes (black) and Ail in liposomes (red), acquired without DNP are shown for comparison and were described previously *(5)*. The yellow rectangle marks LPS-related correlations.

**Figure 2.**
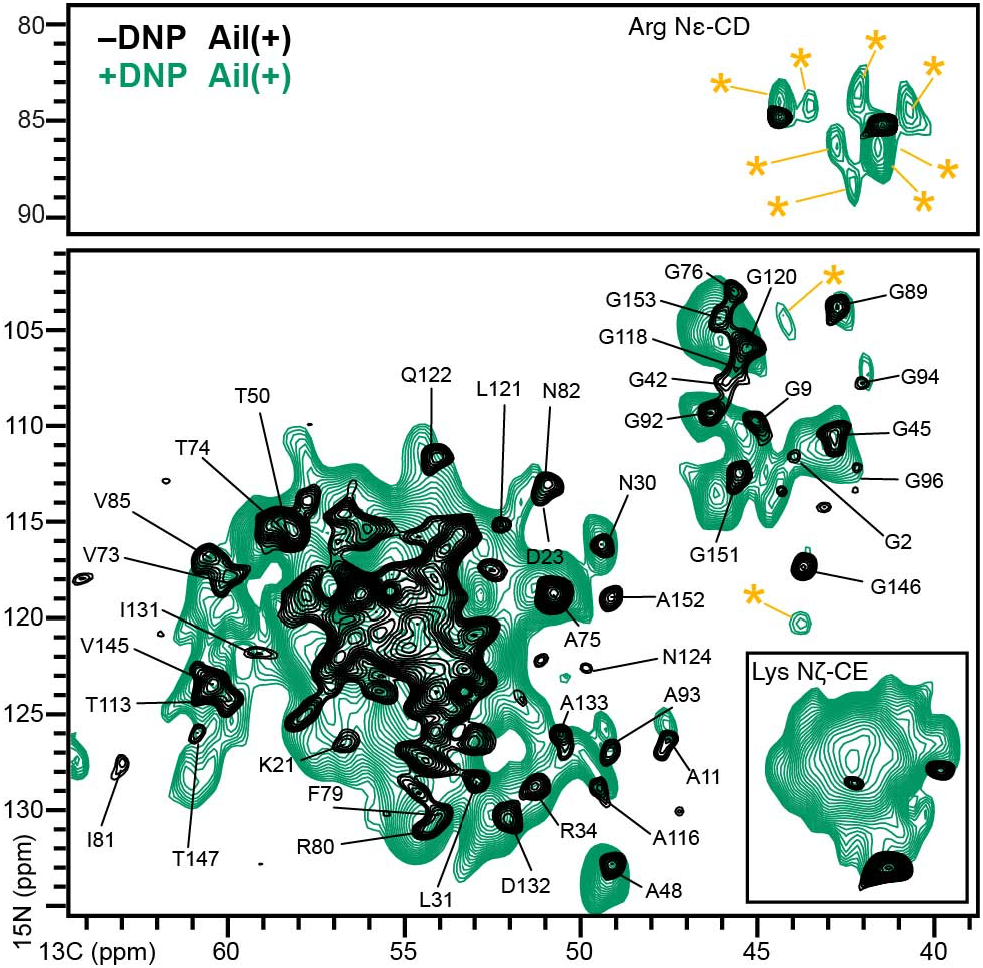
Two-dimensional NCA spectrum of ^15^N,^13^C-Ail in *E. coli* cell envelopes acquired with DNP (green). Resolved assigned peaks are marked. Asterisks denote new unassigned peak observed with DNP. The spectrum from Ail in *E. coli* cell envelopes (black) acquired without DNP is shown for comparison and was described previously *(5)*.

The combined effects of DNP and low temperature also yield a significant enhancement of the signals from Arg sidechains (^13^C: 40–45 ppm; ^15^N: 80–90 ppm) which are observable in the NCA spectra due to the similarity of the Nε-Cδ and N-Cα spin groups (Fig. 2). Eight Arg Nε-Cδ signals are visible, in line with the eight Arg residues in the Ail sequence. Of these, three are buried in the barrel interior near the intracellular membrane surface (R34, R80 and R155), while the others are located at the extracellular membrane surface (R14, R27, R51, R52, R110) where they can interact with LPS polar groups. Here too, the observation of discrete signals at low temperature indicates that their corresponding Arg sites must each adopt a relatively ordered conformational state, since the freezing out of multiple disordered conformational states for each site would be expected to result in extensive line broadening and loss of signal intensity.

To examine the potential of Ail-LPS contacts as a source of conformational order, we acquired two-dimensional NHHC spectra *(32)* where through-space contacts between ^1^H bound to ^13^C and ^15^N nuclei are observed as correlations between their respective ^13^C and ^15^N nuclei (Fig. 3A). We recorded spectra with DNP, at 100K, for Ail(+) and control Ail(–) cell envelopes. Overall, in the Ail(–) cell envelope sample, signals decrease for the amide nitrogen (^15^N: ∼120 ppm) and Arg sidechain regions (^15^N: ∼70 - 80 ppm), while the signals for the Lys side-chains remain essentially the same (^15^N: ∼30 ppm). The dramatic difference in signal intensity for the Lys and Arg regions can also be seen in one dimensional ^13^C slices extracted from the spectra (Fig. 3B, C). A closer inspection of the Arg region reveals that much of this signal reflects intra-residue contacts along the sidechain (Cα, ∼55 ppm; Cβ, 32 ppm; Cγ, 28 ppm; Cδ, 44 ppm). However, the ^13^C signals observed between 62-78 ppm are consistent with the chemical shifts of the peptidoglycan layer and the lipid A portion (glucosamine, 55-100 ppm; carbons from the acyl chains, 25-70 ppm) or the core sugars of LPS *(33)*. The unique presence of these correlations in the Ail(+) sample strongly suggests that they reflect interactions between Arg residues on the protein and the outer leaflet LPS. On the other hand, the negligible difference for the characteristic Lys Nη signals between the Ail(+) and control Ail(–) suggests that interactions between Lys sidechains and LPS are not likely. This is also consistent with our previous solution NMR and MD simulation data, which suggest key roles for Arg side-chains in establishing specific hydrogen bonds with LPS *(23)*. Finally, the decrease in the amide nitrogen signals, which is uniform throughout the whole chemical shift range, most likely reflects the reduced protein content of the control Ail(–) samples.

**Figure 3.**
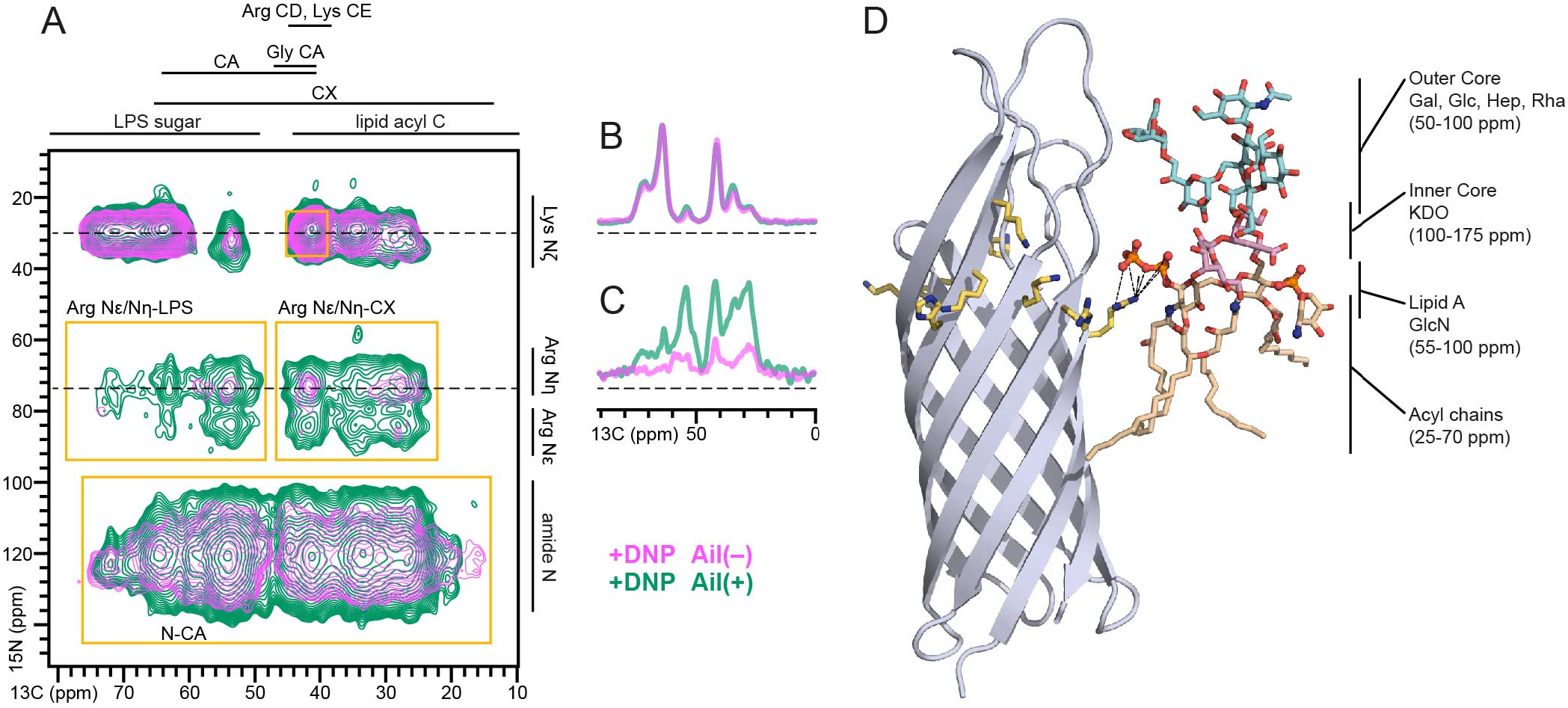
Two-dimensional NHHC spectra acquired with DNP. **(A-C)** Spectra were acquired for (^15^N,^13^C)-Ail(+) (green) or (^15^N,^13^C)-Ail(–) (pink) *E. coli* cell envelopes. Spectral regions of correlations are marked (gold boxes). One-dimensional slices (B, C) were taken at specific ^15^N chemical shifts (dashed lines) of sidechain N from Arg and Lys. **(D)** Snapshot of Ail obtained after 1.5 μs of MD simulation in a *Y. pestis* outer membrane showing that Arg27 in the LPS-recognition motif establishes multiple interactions with lipid A of an LPS molecule. MD simulations were described previously *(23)*.

In summary, we have demonstrated that DNP-enhanced NMR spectroscopy, together with selective labeling strategies, can be used to characterize proteins in their native membrane context with high efficiency and specificity. Despite the line broadening that typically accompanies protein spectra at low temperatures, the spectra of the Ail outer membrane protein are of sufficient quality to allow the identification of many resolved cross peaks and the observation of new peaks that are absent in room temperature spectra. Of note, the new peaks that appear in the Gly and Arg regions of the spectra, for example, display narrow linewidths, suggesting that these residues have well defined conformations at low temperature. The sensitivity enhancement afforded by DNP, on the other hand, has allowed us to observe potential interactions between the Ail Arg e sidechains and LPS of the outer membrane layer, indicating that these residues can play a fundamental role in the restructuring of the lipid environment around the protein. These results showcase the ability of DNP-enhanced NMR spectroscopy to capture elusive interactions in native samples and to expand the size and complexity of biological systems that can be characterized at atomic resolution. We look forward to future studies that extend these capabilities to Ail in whole cells, and to illuminating its interactions with relevant human host proteins.

## ASSOCIATED CONTENT

Methods are provided in the Supporting Information.

## AUTHOR CONTRIBUTIONS

The manuscript was written through contributions of all authors. All authors have given approval to the final version of the manuscript.

## NOTES

The authors declare no competing financial interest.

## ACKNOWLEDGEMENTS

The authors thank Marassi and Debelouchina lab group members for helpful discussions.

