## Supporting Information for "Dynamic nuclear polarization illuminates key protein-lipid interactions in the native bacterial cell envelope"

### MATERIALS AND METHODS

**Protein Expression.** *E. coli* cell envelopes enriched in $^{13}C/^{15}N$ labeled Ail were prepared as described previously *(1)*. Briefly, a plasmid harboring the gene encoding the sequences of Ail and the signal peptide of pectate lyase B was transformed into *E. coli* Lemo21(DE3) cells. Transformed cells were grown to $OD_{600}$~0.4, at 30°C, with vigorous shaking, in M9 minimal media supplemented with Basal Medium Eagle vitamin solution (1% by vol.), ampicillin (100 µg/ml), and chloramphenicol (50 µg/mL). The cells were harvested by low-speed centrifugation (5,000 g , 4°C, 20 min), resuspended in fresh supplemented M9 media, and induced by adding 0.4 mM isopropyl β-D-1-thiogalactopyranoside (IPTG). After reducing the shaking speed and the temperature to 25°C, the cells were cultured for 20 min before adding rifampicin (100 µg/mL), and then for an additional 20 hours, in the dark. Finally, the cells were harvested by low-speed centrifugation, and resuspended in HEPES buffer (10 mM, pH 7.4). To obtain uniformly $^{15}N,^{13}C$ labeled Ail, the cells were grown in unlabeled M9 media and only transferred to $^{15}N,^{13}C$ labeled M9, prepared with $(^{15}NH_4)_2SO_4$ (1 g/L) and $^{13}C_6$-glucose (5 g/L), before induction with IPTG. Control Ail(–) samples were prepared analogously except that they were not transformed with the Ail containing plasmid.

**Isolation of bacterial cell envelopes**. Ail-expressing cells were lysed by three passes through a French Press. After removing cellular debris by centrifugation (20,000 g, 4°C, 1 h), the total cell envelope fraction, including inner and outer membranes, was harvested by ultracentrifugation (100,000 g, 4°C, 1 h), washed three times with sodium phosphate buffer (20 mM, pH 6.5), and then collected by ultracentrifugation. The isolated materials were analyzed by polyacrylamide gel electrophoresis (PAGE) in sodium dodecyl sulfate (SDS), and immuno-blotting with the α-Ail-EL2 antibody *(2)*.

**DNP NMR sample preparation.** For solid-state NMR DNP experiments at cryogenic conditions, a cryoprotectant consisting of glycerol-$d_8$, $D_2O$, and $H_2O$ was added (60/35/5 volume ratio) to the isolated cell envelopes to prevent sample damage during freezing. AMUPol (10 mM final) was added to this mixture, and the sample was packed into a 1.9 mm MAS rotor using low-speed centrifugation.

**DNP NMR Spectroscopy.** Solid-state NMR experiments were performed on a 14.1 T Bruker DNP spectrometer equipped with a NEO console, a 395 GHz gyrotron, and a 1.9 mm $^{1}H/^{13}C/^{15}N$ low temperature MAS probe. Experiments were acquired with a sample temperature of 100 K and a spinning rate of 12,000 Hz. Typical $\pi/2$ pulse lengths were 2.5 µs for $^{1}H$, 3 µs for $^{13}C$, and 7 µs for $^{15}N$. The DNP enhancement was measured by acquiring one-dimensional spectra with or without microwave irradiation at 125 mA power. One and two-dimensional $^{15}N/^{13}C$ NCA spectra were acquired with contact times of 750 µs for $^{1}H/^{15}N$ CP, and 4.0 ms for $^{1}H/^{15}N/^{13}C$ CP. Two-dimensional spectra were acquired with 20 ms $^{13}C$-$^{13}C$ mixing for PDSD, and 400 µs of $^{1}H$-$^{1}H$ mixing for NHHC. The PDSD experiment was collected with 16 scans and 512 points in the indirect dimension. The NCA and NHHC experiments were acquired with 720 scans and 72 points in the indirect dimension each. All experiments used a recycle delay of 5 s.
